# Accurate Automated Segmentation of Autophagic Bodies in Yeast Vacuoles Using Cellpose 2.0

**DOI:** 10.1101/2023.10.23.563617

**Authors:** Emily C. Marron, Jonathan Backues, Andrew M. Ross, Steven K. Backues

## Abstract

Segmenting autophagic bodies in yeast TEM images is a key technique for measuring changes in autophagosome size and number in order to better understand autophagy. Manual segmentation of these images can be very time consuming, particularly since hundreds of images are needed for accurate measurements. Here we describe a validated Cellpose 2.0 model that can segment these images with accuracy comparable to that of human experts. This model can be used for fully automated segmentation, eliminating the need for manual body outlining, or for model-assisted segmentation, which allows human oversight but is still five times as fast as the current manual method. The model and instructions for its use are presented here for the autophagy community.

## Introduction

When an autophagosome fuses with the yeast vacuole, its inner membrane and contents are delivered into the vacuole lumen, forming an autophagic body (APB). Normally, APBs are rapidly degraded by vacuolar hydrolases. However, the inhibition of protease activity in the vacuole, for example by genetic removal of the activating protease Pep4, allows APBs to accumulate so they can be measured^1^. The size and number of APBs in a vacuole corresponds to the size and number of autophagosomes that were formed. Therefore, measuring APBs in different conditions or different mutants can provide valuable insights into the process of autophagosome formation^2–6^.

The only microscopy technique with a high enough resolution to allow measurement of APB size is transmission electron microscopy (TEM). However, since regular TEM visualizes only a thin slice of the cell, many hundreds of cells must be imaged to provide enough data for accurate estimation of original body size and number^7^. Manually labeling the APBs in these images is a laborious process that takes many hours of expert time, creating a clear need for an automated solution.

Rapid advances in computer vision over the past decade have led to the hope that an algorithm could be developed to automatically recognize and label APBs in TEM images. However, this has proven to be a particularly difficult task. Structures in TEM images, unlike those in many fluorescent images, are identified by subtle differences in contrast and texture. Regular TEM images show only a slice of the cell, so there is no three dimensional data to aid in identifying complete structures. (Collecting 3D TEM images through techniques such as serial sectioning is possible but challenging, and not practical for the large sample sizes we need to accurately assess average APB size and number). Finally, APBs can have other cellular membranes inside them that can look like the boundary of a body, while at the same time imperfect TEM preservation can make the actual boundaries incomplete. Together these factors can make it challenging to distinguish one body from the next, and even experts vary somewhat in their labeling of a given image.

Until recently, available tools (including ImageJ, Cellprofiler, Ilastik Multicut, Cellpose 1.0 and a proprietary commercial solution) were woefully inadequate to the task, giving APB segmentations that only remotely resembled the ground truth. However, this changed with the recent release of Cellpose 2.0^8^. Cellpose 2.0 contained two key improvements on Cellpose 1.0 that proved critical for its success in this case: First, it came with a model zoo of pretrained models to speed the training of problem-specific models. Secondly, it had an improved GUI (graphical user interface) that allowed easier human modification of automatically generated labels, making model-assisted labeling feasible^8^. Even though Cellpose 2.0 was primarily designed for labeling of fluorescent images, we decided to try training it on TEM images of APBs, with impressive results – we have been able to train a model that labels ABPs with accuracy comparable to that of human experts.

## Results

Traditional machine learning methods require training on a large number of images to reach optimal performance. Cellpose 2.0 bootstraps this process with the use of pretrained models, and this has previously allowed it to achieve optimal training on specific fluorescent microscopy segmentation problems using only 500-1000 labeled regions of interest (ROIs)^8^. In previously reported Cellpose 2.0 applications, an ROI is typically a cell or nucleus, while in our context an ROI is an APB section. The images we segment typically contain between one and 40 APB sections per image, averaging around 10, raising the possibility that as few as 50-100 images might be sufficient for training. To determine if this estimate would apply to recognizing APBs in electron microscopy images, we trained four separate models using different numbers of labeled APBs. We also trained each model for different numbers of epochs (1 to 999) to determine how many were required for optimal performance; using too many training epochs runs the risk of overtraining the model, which can reduce its performance on novel images. After training, each model was tested on a set of 51 images, and the model’s automated labels were compared to the human-labeled ground truth using the same calculation that Cellpose uses^8,9^: the average precision (AP) at intersection over union (IoU) thresholds of 0.5 and 0.75. An IoU of 0.5 means that 50% of the total pixels of both labels of the same object were in agreement with each other, and indicates that the human and machine had identified roughly the same structure, while an IoU of 0.75 is a “good” agreement between two labels of the same structure. Therefore, if a model had an AP score of 0.65 at IoU threshold = 0.5 and 0.45 at IoU threshold = 0.75, that means it labeled 45% of the APBs the same as the reference segmentation, and labeled a total of 65% fairly similarly.

Figure 1A shows how the accuracy of the models changed during training. We found that each model’s accuracy reached a plateau by about 100 epochs, and evidence of overtraining started around 300 epochs. In addition, we found that increasing the number of samples or epochs did not significantly enhance the model’s performance: Model 2, trained on 1400 APBs, performed only slightly better than Model 0, trained on 350; Model 3, trained on 1837 APBs, actually performed worse than Model 2 (Figure 1A). This is consistent with previous claims that Cellpose 2 requires relatively few ROIs for training, and is beneficial should other researchers desire to train their own models. We selected Model 2 at 201 epochs as our best performing model for further testing and use.

**Figure 1.**
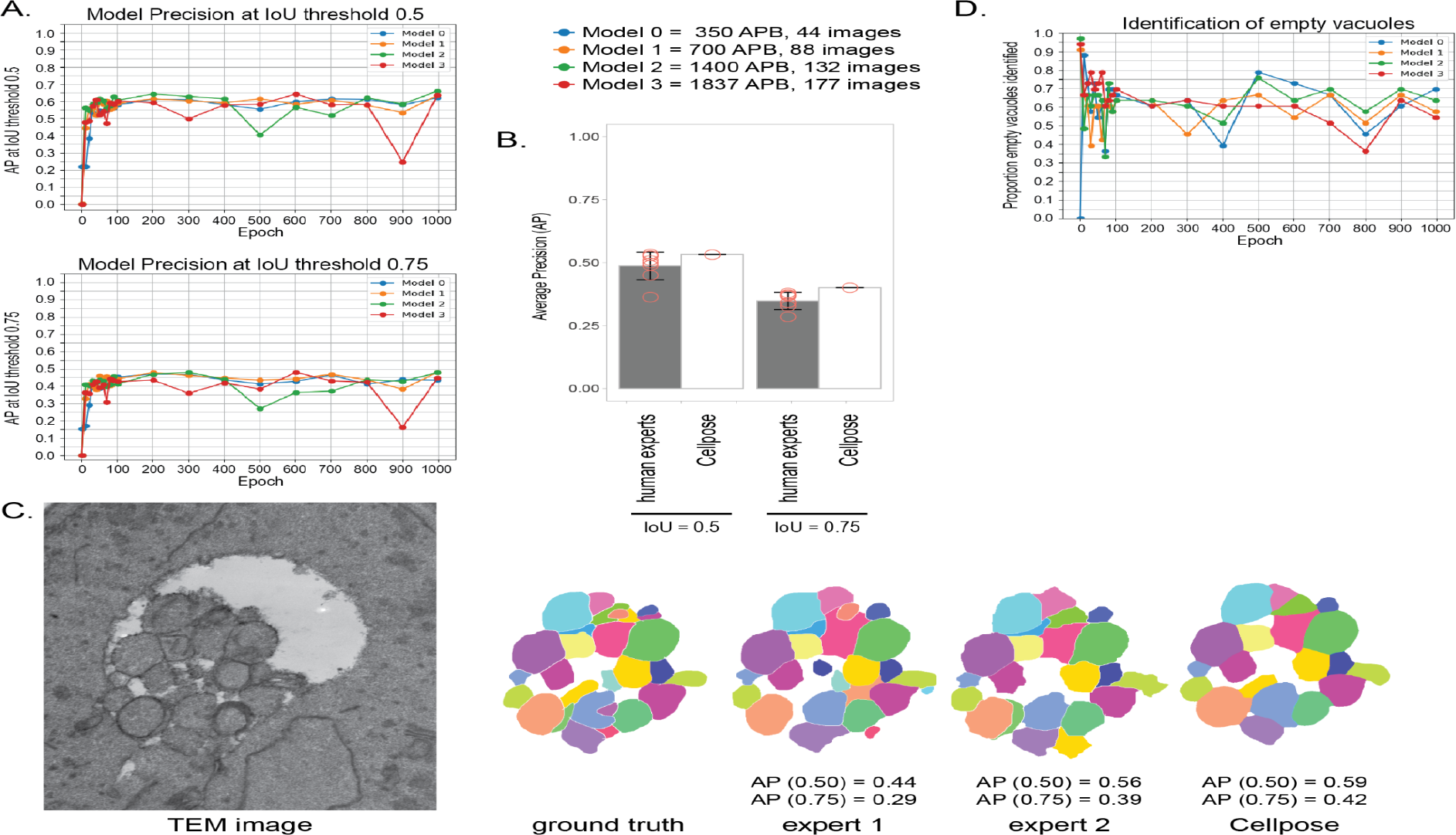
Evaluation of a Cellpose 2 model trained to accurately predict APBs. **(A**) Performance of Cellpose models trained on increasing numbers of APBs for various epochs when tested on 51 images not in the training set, as measured by the average precision at IoU (Intersection over Union) thresholds of 0.5 and 0.75. No additional improvement in performance was seen after 200 epochs of training, and very little improvement was seen beyond 350 APBs, showing that Cellpose 2 can be accurately trained on relatively little data. (**B**) Comparison of the best Cellpose model (Model 2, 201 epochs) to the ABP segmentation performance of eight human experts on nine challenging TEM images. All segmentations were compared to the “ground truth” segmentation of a ninth human expert who had overseen the ground truth segmentation of the training images, and AP values were calculated at IoU thresholds of 0.5 and 0.75. Hollow circles represent the performance of individual segmenters, while bar graphs are the average performance and error bars indicate the standard deviation. Cellpose performed as well as any of the human experts on this task. (**C**) A portion of one of the nine TEM images and examples of various segmentations of that image, including the ground truth reference segmentation, segmentations by two separate human experts, and the segmentation generated by Cellpose using Model 2. AP values relative to the ground truth segmentation are shown. (**D**) The ability of Cellpose models to correctly identify empty vacuoles was tested on 33 images with no APBs; Cellpose’s performance was relatively poor on this task and did not improve with training, likely because it was not possible to include empty vacuole images in the training set.

Model 2 had an AP score of 0.65 at IoU threshold = 0.5. In contrast, published Cellpose 2.0 models for fluorescent or DIC images were able to reach AP scores of 0.7 to 0.8 at IoU threshold = 0.5^8^. One explanation for this lower score would be if TEM images of APBs were particularly difficult to segment even for humans. To test this, we measured between-human agreement on APB labeling by asking eight human experts to segment nine challenging test images. We found the average human AP score to be only 0.49 at IoU threshold = 0.5 and 0.35 at IoU threshold = 0.75 (Figure 1B). This underscores the fact that segmentation of APBs in these images is somewhat ambiguous, and individual scientists may interpret them differently. We used Model 2 to segment those same nine test images, and it achieved an AP score of 0.53 at IoU threshold = 0.5 and AP of 0.40 at IoU threshold = 0.75, suggesting that it can perform this segmentation on par with human experts (Figure 1B,C).

Even though Cellpose 2.0 performs at least as well as a typical expert, a given researcher might want to review Cellpose’s segmentations to make sure they conform to their own best judgment. Fortunately, this is also easy to do, thanks to Cellpose 2.0’s GUI that allows rapid editing of the segmentation. To estimate how much time could be saved by this sort of “model-assisted segmentation,” we used Cellpose 2.0’s GUI with Model 2 loaded to segment 20 new test images one at a time, manually editing any segmentations we were not completely satisfied with. That took a total of 12 minutes of computation time plus 7 minutes of human time for the editing. In contrast, manually segmenting the same 20 images took 39 minutes. Therefore, model-assisted segmentation took less than one fifth as much human time as fully manual segmentation – a major speed improvement.

The only limitation that we found to Cellpose 2.0’s performance is that it was not very good at recognizing images that had no APBs in the vacuole (important for estimation of APB number). It was able to correctly identify only about 70% of empty vacuoles, placing spurious APBs in the others, and this performance did not improve at all with training (Figure 1D). This is likely because we could not include any images with 0 APB in the training data, as the neural network is not set up to train on 0 APB/ROI. It is beyond the scope of this work to make any major changes to the Cellpose 2.0 code. Fortunately, this limitation does not significantly impede the usefulness of Cellpose 2.0 for segmentation of APBs because empty vacuoles can be quickly and easily recognized by humans and require no segmentation, so they can simply be separated from the images to segment before application of Cellpose.

## Discussion

Manual segmentation of large numbers of TEM images has been a rate-limiting factor in the analysis of how different mutations affect autophagosome size and number, as is true of many TEM-based techniques. This not only slows the pace of research but also limits the practical sample size, potentially reducing the accuracy of the final results. In addition, individual variations in segmentation style may lead to issues when comparing results between researchers. Fortunately, advances in machine learning have now made it possible for a computer algorithm to accurately segment ABPs in TEM images. The Cellpose 2.0 model that we present here is as accurate as an independent human expert, but much faster and with reproducible results.

In addition, this model can be further customized as needed. The Cellpose 2.0 GUI allows a user to easily edit the automatically generated segmentations. Moreover, those edited segmentations can be used to further train the model to adjust its style, a procedure described by the Cellpose authors as “human-in-the-loop” training. Since the model requires only a few hundred ROI’s to reach essentially maximum performance, it is even feasible to train an entirely new model if desired. However, using the model presented here without further training or editing of the results would have the advantage of allowing greater reproducibility between studies.

### How to measure APBs using Cellpose 2.0 for labeling

1. Acquire TEM images of APB in yeast vacuoles, carefully following the steps described in section 2.1 of Backues et al. 2014 ^7^
2. Manually sort out images of vacuoles that do not contain any bodies. These will still be used for the analysis of APB number, but do not need to be segmented.
3. Download the trained APB model from https://osf.io/wrez4/
4. Download, install and run Cellpose 2.0 to segment the images containing APBs
  a. For simplest use, install the Cellpose 2.0 GUI using instructions from https://github.com/MouseLand/cellpose. The Cellpose 2.0 GUI allows images to be segmented one at a time: Load the APB model, load the desired image, choose “run model” to generate masks, and then save these as png/tif.
  b. For those with more computational experience, automated segmentation can be performed faster in batch using Pycharm and a Docker container (biocontainers/cellpose:2.1.1_cv2); utilities for this can be found at https://github.com/StevenBackues/cellpose_APB.
5. Copy the mask files to their own folder, and use these to measure the area of each labeled APB and the number of APBs per vacuole. This can be easily done in python using code provided in the utilities or in ImageJ using the MorphoLibJ library^10,11^.
6. Estimate the original size and number distribution of the bodies from this data following the instructions in section 2.3 of Backues et al. 2014^7^.

## Methods

Images of APBs used for the training and test sets were obtained and segmented as previously described^7^. In brief, yeast were starved for 3 hours in media without nitrogen (SD-N) to induce autophagy, and prepared for TEM via chemical fixation. Each image was captured at a magnification of 30,000x, yielding images of 2240 x 2240 pixels, with a scale of 1 px = 2.16 nm. Each image included one yeast near the center, and some had small portions of other yeast near the edges or corners, but no image showed two entire yeast vacuoles. Most images were of yeast with wild-type-sized bodies, but ∼1/4 of images were of a genetic background that produced ∼20% smaller average APB cross-sections (due to reduced ATG7 levels)^6^ to capture this variation. Ground truth manual segmentations were performed by J., S. or R. Backues and verified by S. Backues before use. Training and test data, including images and ground truth segmentations, have been made publicly available at https://osf.io/tuhwn/

For stand-alone use via the GUI, Cellpose 2.0 was downloaded from its official Github repository, https://github.com/mouseland/cellpose. For batch labeling, model training and testing, we used Pycharm Professional and Docker Desktop with the Cellpose 2.0 Docker container biocontainers/cellpose:2.1.1_cv2. All training and tests were run using an nVidia RTX 4090 24GB graphics card; the model can also be used on other systems (GPU or CPU) with essentially identical results. Details of code used can be found at https://github.com/StevenBackues/cellpose_APB.

Our preliminary tests with the pretrained models in Cellpose 2’s model zoo indicated that “CPx” with a diameter parameter of 130 pixels gave the most promising initial results, though not yet with usable accuracy (AP of 0.124 at IoU threshold = 0.5). We thus proceeded to train “CPx” into custom models using different sized sets of APB training data (350, 700, 1400 and 1837 ROIs) and training for 1 to 1000 epochs. Stratification was used to ensure a diverse number of APBs per image (but always more than 0) in both the test and training sets. In addition, we used an empty-vacuole image test set which contained 33 images with no APB. Average Precision (AP) at different IoU thresholds was measured using Cellpose’s built-in testing utility^9^, with AP scores averaged over all 51 test images. The proportion of empty vacuoles correctly identified was calculated using a simple Boolean where True corresponds to correctly predicting that an image had no APB, and False corresponds to any failure to do so (i.e. predicting the presence of any APB in the empty vacuoles). The number of correct predictions was divided by the total number of empty vacuoles tested (33) to give the final score.

## Abbreviations

APB: Autophagic Body
AP: Average Precision
GUI: Graphical User Interface
IoU: Intersection over Union
ROI: Region Of Interest
TEM: Transmission Electron Microscopy

## Acknowledgements

Many thanks to Hayley Cawthon, Ronith Chakraborty, Elizabeth Delorme-Axford, Payton Dunning, Yuchen Feng, Jacquelyn Roberts, Patrick Wall and Zhiping Xie for serving as the human experts for comparing APB segmentation variability. Thanks to Rebecca Backues for performing manual segmentation of some of the images used for training, and to Mark Backues for helpful discussions in the early stages of this project. This work was supported by NSF award #2243163 (RUI: Tools and Approaches for Investigating the Basic Mechanisms of Autophagy) to S. K. Backues, and by Eastern Michigan University.

